# Differential effects of postpartum sleep restriction on maternal and offspring immunity in the rat

**DOI:** 10.1101/2025.09.30.679574

**Authors:** Florencia Peña, Claudio Rodríguez-Camejo, Ana Hernández, Mayda Rivas, Anderson Saravia, Diego Serantes, Juan Pedro Castro, Pablo Torterolo, Teresa Freire, Luciana Benedetto

## Abstract

**Background:** sleep disturbances can trigger a wide range of physiological consequences, affecting hormonal regulation, metabolism, cognitive function, and immune responses. Human mothers worldwide frequently experience sleep restriction and fragmentation, a pattern also observed in other mammalian mothers, such as rats. These alterations may add to sleep disturbances unrelated to motherhood. Considering this, we wondered about the impact of sleep restriction in postpartum mother rats on their immunological status. Furthermore, given that early-life experiences can shape the immune system and that even subtle parental changes can influence offspring development, we hypothesized that maternal sleep loss might also exert detrimental effects on the pups. In this study, we investigated the effects of acute and chronic maternal sleep restriction during the postpartum period on immune parameters in both mother rats and their offspring by analyzing antibody titers and systemic inflammation.

**Methods:** mother rats were surgically implanted with electrodes for polysomnographic recordings and for sleep deprivation (deep electrodes targeting the mesopontine wake-promoting area). From postpartum day 5 to day 9, lactating dams were randomly assigned to one of three groups: chronic sleep restriction (CSR; 6 h of sleep deprivation per day for five consecutive days), acute sleep restriction (ASR; 6 h of sleep deprivation only on postpartum day 9), or control (undisturbed). On postpartum day 9, mothers were milked, and blood samples from both mothers and pups were subsequently collected. ELISA assays quantified IL-17A, IL-6, IgG, and IgG2a in maternal serum; IgG and IgG2a in milk; and IgG in pup serum. Hematological parameters, including leukocyte profiles, were also assessed in peripheral blood of dams and pups.

**Results:** maternal immune parameters analyzed remained unaffected by sleep restriction. IgG levels were lower in male pups from mothers subjected to ASR (5560 ± 734 µg/mL) compared with the control group (8666 ± 463 µg/mL; p = 0.025), whereas female pups showed no significant changes. Additionally, both female (4.10 ± 0.58) and male (3.81 ± 0.42) pups from dams subjected to CSR exhibited higher absolute lymphocytes counts relative to the control group (females: 2.28 ± 0.25, p = 0.004; males: 2.44 ± 0.25; p = 0.029).

**Conclusions:** Chronic and acute maternal sleep restriction had distinct impacts on offspring immunity, altering serum antibody and leukocyte profiles, while leaving maternal parameters unaffected. These results indicate that maternal sleep loss can influence the offspring even in the absence of detectable maternal immune alterations, with male pups being especially susceptible.

## 1. Introduction

Sleep is a vital process for most species. Periods of sleep deprivation or restriction have been shown to alter numerous biological functions, including hormonal regulation, cognitive performance, and immune responses to infectious agents ^1–4^. Despite this, sleep deprivation and/or fragmentation are common traits of motherhood worldwide ^5–9^, and a characteristic shared with other mammals studied ^10–15^. Furthermore, additional factors unrelated to maternal care, such as work schedules ^16,17^ or certain health conditions ^18^, may exacerbate the sleep disturbances already present during motherhood ^19^, raising the question of its consequences.

The immune system and sleep are closely related ^4,20–25^. Some proinflammatory cytokines, such as interleukin 1 (IL-1), tumor necrosis factor (TNF-α) and interferon alfa (IFN-α), present somnogenic activity, increasing NREM sleep and the amplitude of the slow waves observed in the electroencephalographic (EEG) recordings, suggesting that cytokines are involved in normal sleep regulation ^4,24,26–28^. In turn, the levels of these cytokines change in response to sleep disturbances. For instance, sleep restriction promotes the secretion of pro-inflammatory cytokines, such as IL-1β, IL-6, IL-17A, and TNF-α ^29–33^, and reduces the production of anti-inflammatory cytokines, such as IL-10 ^4,33^. Besides, humoral mediators of the adaptive response are also affected by sleep deprivation, such as the antibodies. Total sleep deprivation increases the serum concentration of IgG, IgM, and IgA ^34–37^, whereas REM sleep deprivation has been shown to decrease levels of IgG1 and IgG2a ^38^. Moreover, insufficient sleep impairs the humoral response to vaccination, as evidenced by reduced antigen-specific antibody titers ^39,40^, suggesting that sleep loss compromises the adaptive immune system’s ability to mount an effective response following immunization. Finally, sleep also plays an active role in regulating cellular immune function and recirculation ^4,25,41,42^, as sleep restriction has been shown to decrease blood leukocyte counts, tissue migration and phagocytic capacity ^43–49^.

Regarding the postpartum period, little is known about the effects of sleep alterations on immune function. It has been reported that human mothers experiencing higher levels of daytime sleepiness and fatigue, as assessed by questionnaires, correlate with more symptoms of infection and higher levels of IL-1β ^50,51^. Furthermore, IL-6 and IL-10 concentrations do not differ between women with or without insomnia during the postpartum period ^52^. However, sleeping five hours or less per day during the first postpartum year is associated with elevated IL-6 levels ^53^. Similarly, lactating cows that are not allowed to adopt the resting posture necessary for sleep, express higher levels of the inflammatory cytokines TNF-α and IL-6 ^54^. Taken together, these findings suggest that sleep disturbances may also contribute to a proinflammatory state during the postpartum period. Taking this into consideration, we wonder if mother rats subjected to sleep restriction in addition to the intrinsic sleep disturbances of the maternal period would show immune alterations.

Furthermore, even minor environmental or parental behavior changes within the normal range can alter offspring development ^55,56^, including the immune system ^57–59^. As we previously showed, sleep restriction in mother rats induced changes in maternal behavior and in the macronutrient composition of their milk ^60^. Therefore, we wondered whether maternal sleep loss might impact certain immune parameters in the mother–offspring dyad.

In the current study, we analyzed the impact of acute and chronic sleep restriction during postpartum on immune parameters in mother rats and their pups.

## 2. Methods

### 2.1. Animals and housing

A total of 26 Wistar primiparous female rats (250-320 g) and their pups between postpartum days 5 to 9 (PPD5-9, birth = day 0) were included in this study. The present study is a continuation of our previous work ^60^ and was carried out using the same cohort of animals.

A day before delivery, pregnant rats were kept in individual transparent cages (40×30×20 cm) containing shredded paper towels for nest-building, and food and water available *ad libitum*. The animals were placed in a temperature-controlled chamber (22 ± 1 °C) with a 12/12 h light/dark cycle with zeitgeber time (ZT) 0–12 marking the lights-on phase of the day. This chamber was soundproof and electromagnetically shielded, equipped with slip rings and cable connectors for bioelectrical recordings. All the experimental procedures were approved by the Institutional Animal Care Committee (protocol number 070151-000008-22) and were undertaken in conformity with the “Guide to the Care and Use of Laboratory Animals” (8th edition, National Academy Press, Washington D. C., 2011) ^61^.

### 2.2. Electrode implantation and bioelectrical signal acquisition

The surgical procedures were similar to those in previous studies of our group ^60,62–64^. The mother rats were anesthetized with an intraperitoneal injection of a ketamine/xylazine/acepromazine maleate mixture (90/2.8/2.0 mg/kg) on PPD1. Unilateral cortical EEG electrodes were placed in the following cortices: frontal (AP = + 3.0, ML = 2.0; from Bregma), parietal (AP = −4.0, ML = 3.0), occipital (AP = −7.0, ML = 3.0) and cerebellum as a reference (AP = −11.0, ML = 0.0) ^65^. Lastly, we implanted a unilateral bipolar braided nichrome electrode (550 µm distance between the electrode tips) in the pontine peduncle tegmentum (PPT; AP = –7.5, ML = 2.0 and H = 8.0 mm from the skull surface ^65^) for the deep brain electrical stimulation (DBES). In addition, two electrodes were inserted into the neck muscles for electromyogram (EMG) recording, and two stainless-steel screws were fixed into the skull as anchors. The electrodes were soldered to a socket and secured with dental acrylic.

A single dose of ketoprofen (5 mg/kg, s.c.) was administered prior to surgery. At the end of the procedure, rats were injected with 0.9% sterile saline (10 ml/kg; s.c.), a single dose of systemic antibiotic (ceftriaxone 60 mg/kg; i.p.), and a topical antibiotic over the injury (Crema 6A, Labyes). During surgery, the pups were maintained in their home cage under a heat lamp, and before each mother was reunited with her pups, the pups were culled to four males and four females ^66,67^.

Polysomnographic recordings were obtained by amplifying (×1000), filtering (0.1–500 Hz), sampling (1024 Hz, 16-bit), and storing the bioelectrical signals using Spike2 software (Cambridge Electronic Design).

### 2.3. Experimental design and sessions

In brief, all recordings were performed during the light phase of the cycle (from ZT 3 to ZT 9.5) for five days between PPD5-9. All mothers were recorded every day after being randomly assigned to one of the three independent experimental groups: CSR group (sleep deprivation every day, n=8), ASR group (sleep deprivation only on PPD9, n=8) or control group (C; all days recorded and allowed to sleep freely during the entire sessions, n = 9). A summary of the experimental design is shown in Figure 1A.

**Figure 1.**
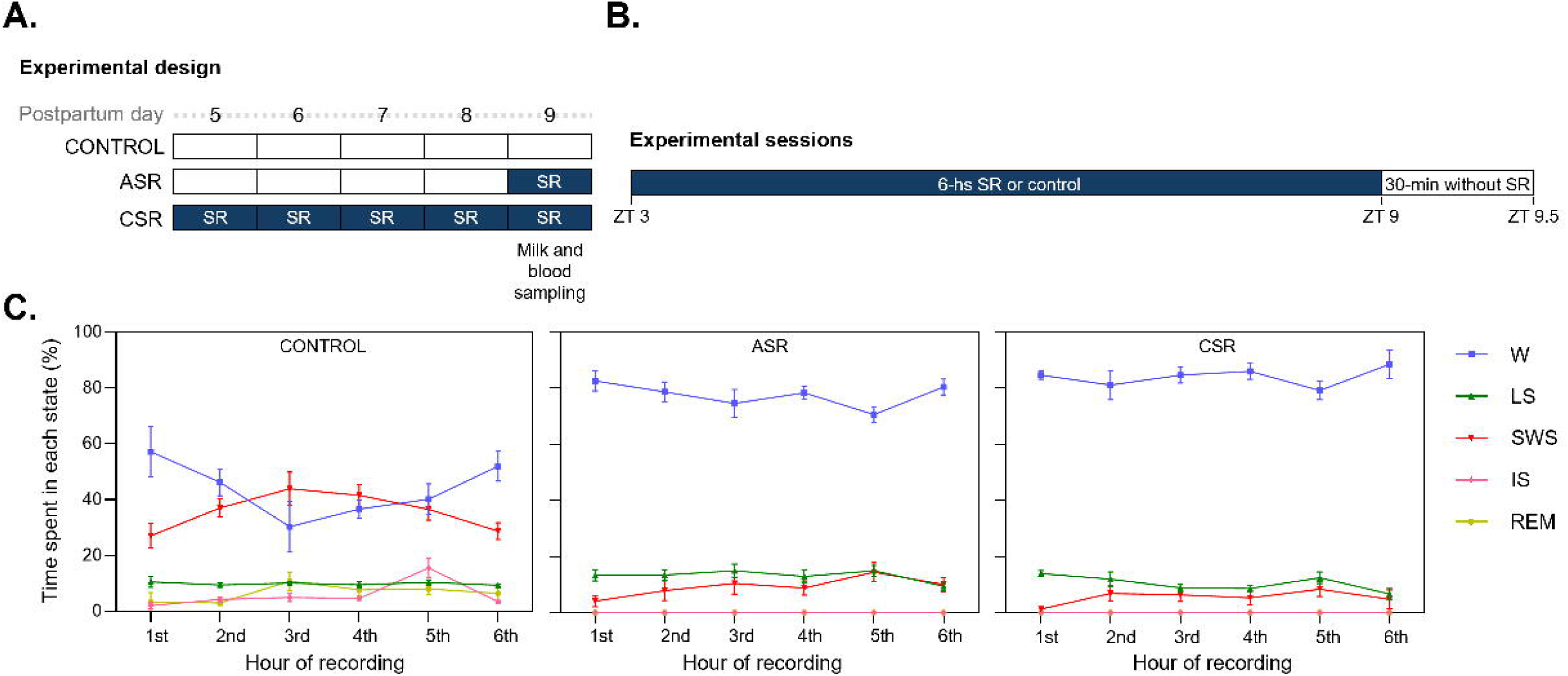
Experimental design and sleep restriction across an experimental session. **A.** The figure shows the experimental design represented by a timeline. The experimental sessions were conducted between postpartum days 5-9 (PPD5-9) and consisted of sleep restriction (SR) or control procedures. We randomly assigned the mother rats to one of the three independent groups: five days of SR (chronic sleep restriction, CSR); one day of SR (acute sleep restriction, ASR) or undisturbed sleep (control). Milk and serum samples were collected only at the end of the PPD9 session. **B.** The figure shows an experimental session, indicating the 6-hour sleep restriction or control recording period, plus the following 30-minute without sleep restriction. **C.** The graphs show the percentage of time spent in each sleep-wake stage during the six-hour sleep restriction recording on PPD9. For more details about the effectiveness of sleep restriction see our previous work ^60^. Data is presented as mean ± standard error from five rats per group. Each state was compared using one-way ANOVA and Tukey test as *post hoc*. ZT: zeitgeber time; W, wakefulness; LS, light sleep; SWS, slow wave sleep; IS, intermediate stage; REM, rapid eye movement sleep.

During each session, a six and a half hour polysomnographic recording was conducted. Depending on the assigned group, the rats either experienced six hours of sleep deprivation or slept freely (Figure 1A). After that, sleep-wake states were recorded for 30 minutes but no sleep restriction was performed in any experimental group (Figure 1B; for more experimental details please see ^60^).

At the end of all experimental sessions (PPD9), dams were anesthetized with a mixture of ketamine/xylazine/acepromazine maleate (90/2.8/2.0 mg/kg, *i.p.*) for milk extraction and subsequently were euthanized for blood collection.

### 2.4. Sleep restriction through deep brain electrical stimulation

The sleep restriction procedures were carried out according to our previous work ^64^. Regarding the sleep deprivation method, we observed that in lactating rats, DBES is as effective as gentle handling in inducing sleep deprivation, while causing less disruption to maternal behavior ^60,64^. For more details about the sleep deprivation method see ^60^.

### 2.5. Milk sampling

The procedure for milk collection was described in our previous work ^60^. Briefly, we covered four nipples of each dam with leukotape to collect milk samples and allow pup feeding simultaneously, thereby avoiding the need for oxytocin injections ^68,69^ or dam-pups separation during sessions ^70^.

The milk was collected in a tube and kept in ice until the end of the extraction. The volume obtained from each mother ranged between 250 μL and 900 μL during a milking period of 40 minutes (for more details see ^60^).

The analysis of immune components in milk was conducted on the aqueous phase, which was separated by centrifuging whole milk for 30 minutes at 15.000 rpm and 4°C, and stored at –80°C until assayed ^60^.

### 2.6. Serum sampling

Immediately after the milking procedure, blood samples from dams were obtained through exsanguination via an intracardiac bleed (2 ± 0.5 mL per rat). Blood samples from the pups were obtained via decapitation (0.5 ± 0.1 mL per pup).

For hematological and leukocyte parameters, an aliquot of the blood from mothers and pups was collected in tubes containing sodium citrate at a final concentration of 4% (1:10 citrate-to-blood ratio). The remaining blood was kept cold (4°C) overnight, centrifuged at 2500 g for 15 minutes at 4°C, and then the serum was extracted, aliquoted and stored at –80°C until analysis.

### 2.7. Analysis

#### 2.7.1. Sleep and wakefulness parameters

For the last experimental sessions (PPD9), wakefulness (W), light sleep (LS), slow wave sleep (SWS), intermediate stage (IS), and REM sleep were classified into 5-second epochs and identified through visual inspection according to standard criteria^62,63,71,72^.

Sleep restriction was effectively carried out using DBES. The differences in total sleep time and the total time spent in each stage, as well as the differences between groups during the last half hour, are reported in our previous work ^60^. Here, we show the hour-to-hour differences in W, LS, SWS, IS and REM across the six-hour sleep restriction or control periods during the last experimental session of all groups (Figure 1C). The percentage of each state did not change throughout the 6 hours within each group. It is noteworthy that SWS was markedly suppressed, whereas IS and REM sleep was completely abolished in both deprivation groups.

#### 2.7.2. Cytokines quantification in maternal serum samples

Maternal serum concentrations of IL-17A and IL-6 were analyzed by sandwich ELISA according to the following specifications. For IL-6 determinations, high-binding 96-well plates (Thermo Scientific™ Nunc™, 442404) were coated with 50 µL per well of a primary antibody specific for IL-6 (BD Biosciences, Cat No. 550644), diluted 1:400 in 100 mM phosphate-buffered saline (PBS) pH 9, and incubated overnight at 4 °C. Plates were then washed three times with 0.1% PBS-Tween. Subsequently, wells were blocked with 100 µL of 1% gelatin in PBS and incubated for 1 hour at 37 °C. After blocking, three additional washes with 0.1% PBS-Tween were performed. Next, the samples and serial dilutions of recombinant IL-6 (BD Biosciences, Cat No. 557008) for the standard curve (ranging from 12.6 to 2000 pg/mL) were incubated for 1 hour at 37 °C, followed by three washes with 0.1% PBS-Tween. Next, a biotin-conjugated secondary antibody against IL-6 (BD, Cat No. 550642), diluted 1:500 in 0.1% PBS-Tween with 1% gelatin, was incubated for 1 hour at 37 °C. After three additional washes with 0.1% PBS-Tween the plates were incubated with streptavidin-peroxidase (BioLegend, Cat No. 405210), diluted 1:1000, for 1 hour at 37 °C, followed by five washes with 0.1% PBS-Tween. For detection, 200 µL of o-phenylenediamine dihydrochloride (OPD, 0.5 mg/mL) with 0.12% H₂O₂ in 0.1 M citrate-phosphate buffer pH 5, was added. The reaction was stopped with 25 µL of 3 M HCl. Absorbance was measured at 492 nm, and IL-6 concentrations were calculated based on the standard curve.

The concentration of IL-17A was determined using a commercial ELISA kit (MAX™ Deluxe Set Rat IL-17A, BioLegend, Cat No. 437904) following the manufacturer’s instructions. Results were calculated from a standard curve ranging from 15.6 to 2000 pg/mL, prepared in its corresponding assay diluent.

No sample dilutions were performed for any of the cytokine analyses.

#### 2.7.3. Antibody analysis in serum and milk samples

The concentrations of IgG were analyzed in maternal and pups’ serum, and in the aqueous phase of maternal milk, while IgG2a was measured only in maternal samples due to the volume availability of pups’ serum.

In the case of the pups, total IgG was measured using pooled serum samples separated by sex, comprising four females and four males per dam. These measurements were performed using in-house ELISA methods previously standardized in our laboratory ^73^. The assays were conducted according to the following specifications:

High-binding 96-well plates (Greiner Bio-One, Frickenhausen, Germany) were coated overnight at 4°C with 100 µL per well of the appropriate capture antibody: goat anti-rat IgG (R5130, Sigma-Aldrich, St. Louis, MO) or mouse anti-rat IgG2a (553918, BD Pharmingen, NJ, USA), diluted 1:1000 and 1:200 in PBS, respectively. After discarding the contents, plates were washed with 0.05% PBS-Tween-20 and blocked with 200 µL per well of 1% gelatin in PBS for 1 hour at 37°C.

Following additional washes, 100 µL per well of appropriately diluted standards and samples were added and incubated for 2 hours at 37°C in PBS containing 1% gelatin and 0.05% Tween-20. Purified rat IgG and IgG2a (553992, BD Pharmingen) were used to generate standard curves, with serial dilutions starting from 150 ng/mL and 2 µg/mL, respectively.

For detection, 100 µL per well of the appropriate conjugate was incubated for 1 hour at 37°C. To detect total IgG, horseradish peroxidase (HRP)-conjugated rabbit anti-rat IgG (A5795, Sigma-Aldrich) was used at a 1:4000 dilution. For IgG2a detection, a biotinylated mouse monoclonal antibody (553894, BD Pharmingen) was used at a 1:100 dilution, followed by incubation with an HRP-conjugated extravidin solution (ExtrAvidin-HRP, E2886, Sigma-Aldrich) diluted 1:3000.

After washing, the colorimetric reaction was developed by adding 100 µL per well of substrate solution (0.1 M of sodium acetate plus 0.4 mM TMB and 0.004% H_2_O_2_). The reaction was stopped with 1 N H₂SO₄, and optical densities were measured at 450 nm using an ELISA plate spectrophotometer (LabSystems, Multiskan MS).

#### 2.7.4. Hematological parameters and differential leukocyte count

A complete blood count was performed on anticoagulated blood samples to quantify red blood cells and the different populations of white blood cells (lymphocytes, neutrophils, monocytes, eosinophils, and basophils) using an automated hematology analyzer (BC-5000 Vet, Mindray, China). For all experimental groups, the complete blood count was performed on individual maternal and pup samples.

#### 2.7.5. Statistics

Data that did not follow a normal distribution (Shapiro–Wilk test, p < 0.05) are presented as median ± SIQR (semi-interquartile range), while data that followed a normal distribution are presented as mean ± SEM (standard error of the mean). Statistical differences among maternal parameters were evaluated using the Kruskal–Wallis test followed by Dunn’s *post hoc* test for non-normally distributed data, or one-way ANOVA followed by Tukey’s *post hoc* test for normally distributed data. Differences in pups were analyzed using two-way ANOVA, followed by Tukey’s *post hoc* test, with experimental condition (control, ASR, CSR) and sex (female or male) as factors. Spearman correlation matrices and simple linear regressions were performed to evaluate relationships between variables. The criterion used to discard the null hypotheses was p < 0.05. All statistical analyses were performed in GraphPad Prism 10, Dotmatics.

## 3. Results

### 3.1. Humoral immune parameters in dams and pups

No significant differences were observed in IgG levels among the groups in either maternal serum (F = 2.65; p = 0.094), milk (F = 2.13; p = 0.143), or the IgG milk/serum ratio (F = 0.49; p = 0.622; Figure 2A-C). Similarly, no significant group differences were found in IgG2a levels in maternal serum (F = 0.10; p = 0.904), milk (F = 0.17; p = 0.845), or the IgG2a milk/serum ratio (F = 0.51; p = 0.609; Figure 2D-F).

**Figure 2.**
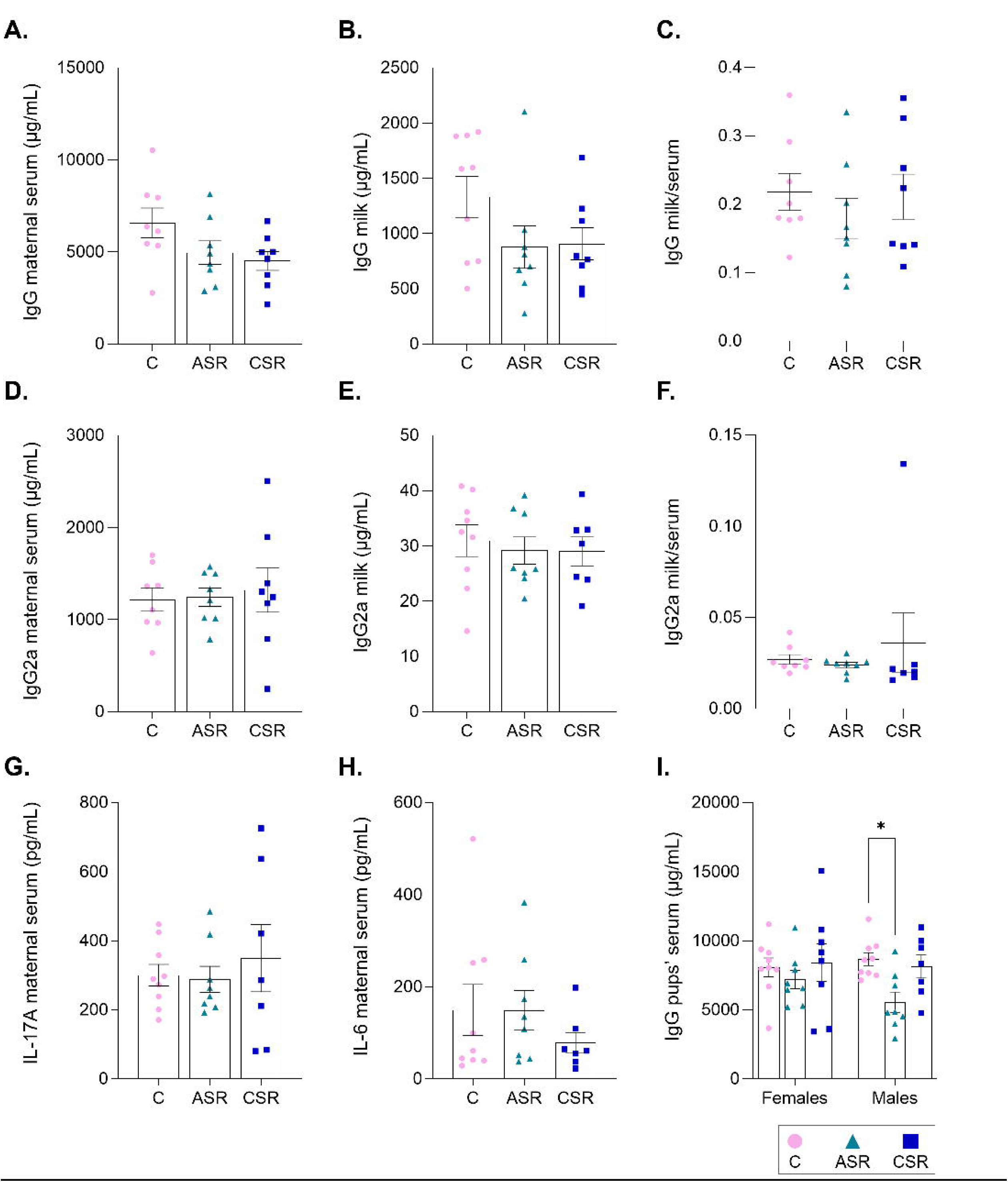
Effects of maternal sleep restriction on humoral immunological parameters in the serum and milk of dams and pups. **A–B.** IgG concentrations in maternal serum **(A)** and breast milk **(B)**. **C.** Ratio between serum and milk IgG concentrations. **D–E.** IgG2a concentrations in maternal serum **(D)** and breast milk **(E)**. **F.** Ratio between serum and milk IgG2a concentrations. **G–H.** IL-17A **(G)** and IL-6 **(H)** concentrations in maternal serum. **I.** IgG concentrations in the serum of female and male pups. All samples were collected on PPD9. For pups, IgG was measured in serum pooled by sex. Data are presented as mean ± standard error. IgG, IgG2a, and IL-17A from maternal data were analyzed using one-way ANOVA followed by Tukey’s *post hoc* test, while IL-6 data were analyzed using the Kruskal–Wallis test followed by Dunn’s *post hoc* test. Pups’ data were analyzed with two-way ANOVA followed by Tukey *post hoc* test. C, control; ASR, acute sleep restriction; CSR, chronic sleep restriction.

There were no statistically significant differences among groups in IL-17A levels (F = 0.30; p = 0.747) or IL-6 concentrations in maternal serum (H = 0.35; p = 0.841; Figure 2G-H).

Regarding the pups, the two-way ANOVA revealed a significant main effect of experimental condition (p = 0.032). Specifically, serum IgG levels were lower in male offspring of ASR dams (5560 ± 734 µg/mL) compared with the control group (8666 ± 463 µg/mL; p = 0.025; Figure 2I). Although the effect was evidenced only in male pups, the two-way ANOVA revealed no significant main effect of sex (p = 0.513) and no significant interaction between sex and experimental condition (p = 0.386).

### 3.2. Hematological parameters of dams and pups

No significant differences were observed in maternal hematological parameters (Table 1). Regarding the pups, two-way ANOVA showed a significant main effect of experimental condition on total leukocyte count (p = 0.001), with females in the CSR group (6.50 ± 1.03) higher than controls (3.61 ± 0.45; p = 0.011) and ASR (4.17 ± 0.62; p = 0.041). Males showed a similar trend (CSR: 5.92 ± 0.85 vs. control: 3.74 ± 0.37; p = 0.059; Table 2). The increase was mainly due to lymphocytes, which also showed a significant experimental condition effect (p = 0.0003), with CSR females values (4.10 ± 0.58) higher than control (2.28 ± 0.25; p = 0.004) and ASR (2.80 ± 0.38; p = 0.040), and CSR males values (3.81 ± 0.42) higher than controls (2.44 ± 0.25; p = 0.029; Table 2).

**Table 1.** Effects of sleep restriction on maternal hematological parameters.

| Parameter | Unit | Control | ASR | CSR | Statistic (F) | p-value |
| --- | --- | --- | --- | --- | --- | --- |
| Leukocytes | 10<sup>3</sup>/μL | 7.28 ± 1.41 | 7.35 ± 0.93 | 6.60 ± 1.07 | 0.12 | 0.887 |
| Neutrophils | 10<sup>3</sup>/μL | 1.89 ± 0.56 | 2.25 ± 0.33 | 1.51 ± 0.36 | 0.66 | 0.530 |
| Lymphocytes | 10<sup>3</sup>/μL | 4.66 ± 0.90 | 4.13 ± 0.57 | 4.16 ± 0.54 | 0.18 | 0.837 |
| Monocytes | 10<sup>3</sup>/μL | 0.56 ± 0.19 | 0.79 ± 0.13 | 0.73 ± 0.22 | 0.45 | 0.644 |
| Eosinophils | 10<sup>3</sup>/μL | 0.15 ± 0.05 | 0.16 ± 0.04 | 0.16 ± 0.03 | 0.02 | 0.979 |
| Basophils | 10<sup>3</sup>/μL | 0.02 ± 0.01 | 0.04 ± 0.01 | 0.02 ± 0.01 | 1.97 | 0.165 |
| Erythrocytes | 10<sup>6</sup>/μL | 4.85 ± 0.21 | 4.81 ± 0.15 | 4.74 ± 0.30 | 0.07 | 0.937 |
| Hematocrit | % | 31.98 ± 1.32 | 32.81 ± 0.90 | 30.88 ± 1.56 | 0.57 | 0.575 |
| Hemoglobin | g/dL | 10.99 ± 0.33 | 11.23 ± 0.24 | 10.77 ± 0.46 | 0.44 | 0.653 |
| MCV | fL | 65.94 ± 2.04 | 68.63 ± 2.29 | 65.93 ± 1.25 | 0.64 | 0.538 |
| MCH | pg | 22.83 ± 0.69 | 23.39 ± 0.60 | 21.88 ± 0.50 | 1.51 | 0.246 |
| MCHC | g/L | 347.5 ± 10.03 | 343.3 ± 5.10 | 335.6 ± 3.83 | 0.70 | 0.507 |
| RDW-CV | % | 23.86 ± 1.87 | 24.98 ± 2.00 | 25.81 ± 1.78 | 0.26 | 0.772 |
| RDW-SD | fL | 67.35 ± 6.05 | 72.89 ± 8.22 | 72.96 ± 6.40 | 0.22 | 0.801 |
| Platelets | 10<sup>3</sup>/μL | 814.0 ± 142.2 | 728.6 ± 101.6 | 677.1 ± 84.87 | 0.37 | 0.700 |
| MPV | fL | 7.31 ± 0.26 | 6.98 ± 0.35 | 7.16 ± 0.32 | 0.30 | 0.747 |
| PDW | % | 17.33 ± 0.38 | 17.03 ± 0.30 | 17.10 ± 0.36 | 0.22 | 0.802 |
| PCT | % | 0.55 ± 0.10 | 0.54 ± 0.09 | 0.49 ± 0.07 | 0.11 | 0.893 |

The results correspond to hematological parameters of dams based on samples collected on postpartum day 9. Data are presented as mean ± SEM and were analyzed using one-way ANOVA followed by Tukey's *post hoc* test. ASR, acute sleep restriction; CSR, chronic sleep restriction; MCV, mean corpuscular volume; MCH, mean corpuscular hemoglobin; MCHC, mean corpuscular hemoglobin concentration; RDW-CV, red blood cell distribution width – coefficient of variation; RDW-SD, red blood cell distribution width – standard deviation; MPV, mean platelet volume; PDW, platelet distribution width; PCT, plateletcrit.

**Table 2.**
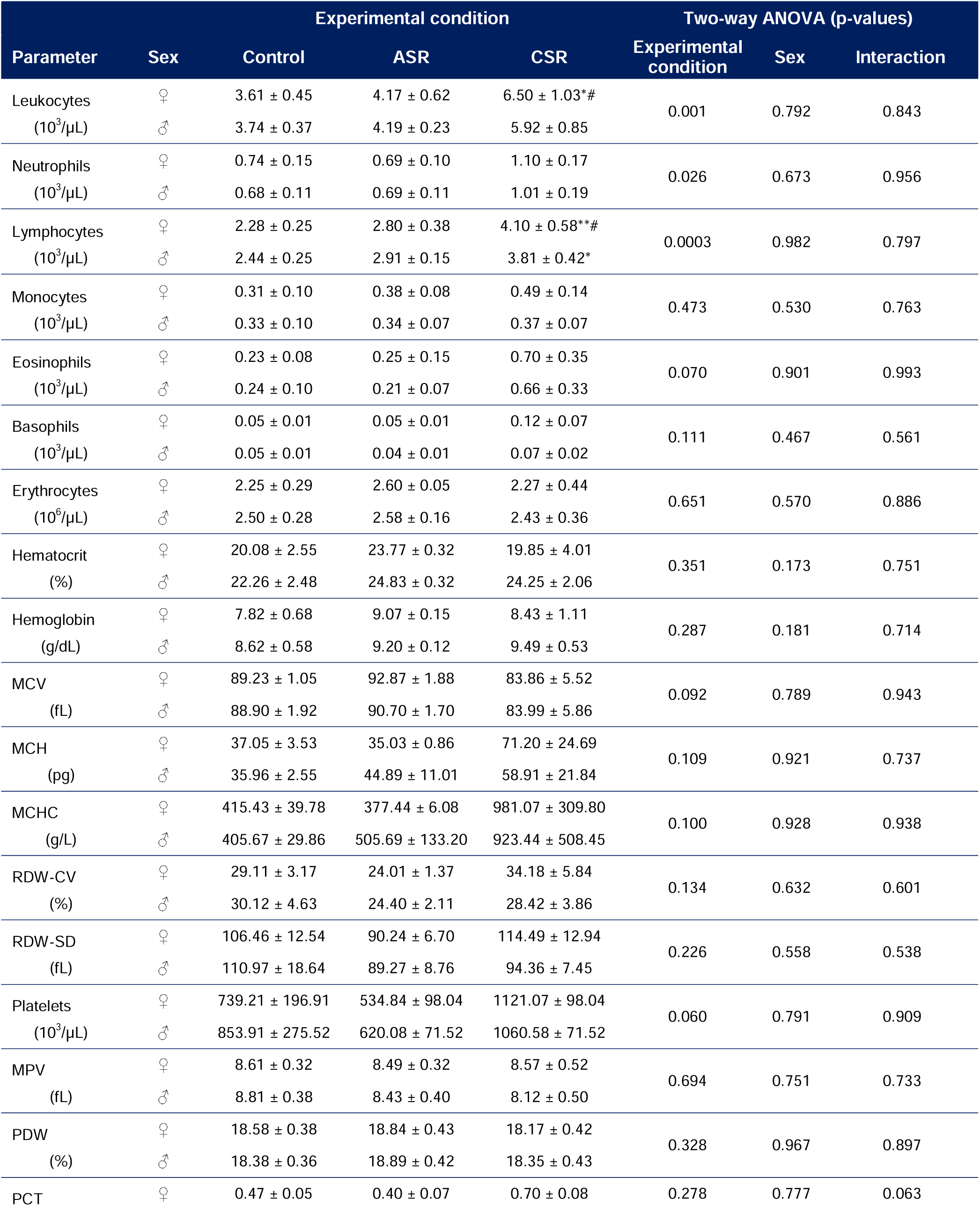
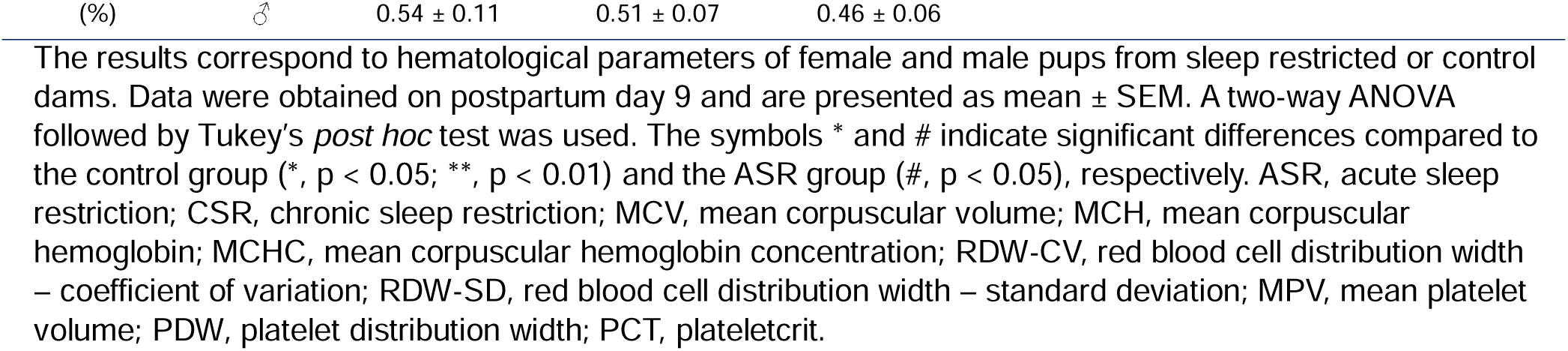
Effects of maternal sleep restriction on female and male pups’ hematological parameters.

| Parameter | Sex | Experimental condition |  |  | Two-way ANOVA (p-values) |  |  |
| --- | --- | --- | --- | --- | --- | --- | --- |
|  |  | Control | ASR | CSR | Experimental condition | Sex | Interaction |
| Leukocytes<br>(10<sup>3</sup>/μL) | ♀ | 3.61 ± 0.45 | 4.17 ± 0.62 | 6.50 ± 1.03*# | 0.001 | 0.792 | 0.843 |
|  | ♂ | 3.74 ± 0.37 | 4.19 ± 0.23 | 5.92 ± 0.85 |  |  |  |
| Neutrophils<br>(10<sup>3</sup>/μL) | ♀ | 0.74 ± 0.15 | 0.69 ± 0.10 | 1.10 ± 0.17 | 0.026 | 0.673 | 0.956 |
|  | ♂ | 0.68 ± 0.11 | 0.69 ± 0.11 | 1.01 ± 0.19 |  |  |  |
| Lymphocytes<br>(10<sup>3</sup>/μL) | ♀ | 2.28 ± 0.25 | 2.80 ± 0.38 | 4.10 ± 0.58**# | 0.0003 | 0.982 | 0.797 |
|  | ♂ | 2.44 ± 0.25 | 2.91 ± 0.15 | 3.81 ± 0.42* |  |  |  |
| Monocytes<br>(10<sup>3</sup>/μL) | ♀ | 0.31 ± 0.10 | 0.38 ± 0.08 | 0.49 ± 0.14 | 0.473 | 0.530 | 0.763 |
|  | ♂ | 0.33 ± 0.10 | 0.34 ± 0.07 | 0.37 ± 0.07 |  |  |  |
| Eosinophils<br>(10<sup>3</sup>/μL) | ♀ | 0.23 ± 0.08 | 0.25 ± 0.15 | 0.70 ± 0.35 | 0.070 | 0.901 | 0.993 |
|  | ♂ | 0.24 ± 0.10 | 0.21 ± 0.07 | 0.66 ± 0.33 |  |  |  |
| Basophils<br>(10<sup>3</sup>/μL) | ♀ | 0.05 ± 0.01 | 0.05 ± 0.01 | 0.12 ± 0.07 | 0.111 | 0.467 | 0.561 |
|  | ♂ | 0.05 ± 0.01 | 0.04 ± 0.01 | 0.07 ± 0.02 |  |  |  |
| Erythrocytes<br>(10<sup>6</sup>/μL) | ♀ | 2.25 ± 0.29 | 2.60 ± 0.05 | 2.27 ± 0.44 | 0.651 | 0.570 | 0.886 |
|  | ♂ | 2.50 ± 0.28 | 2.58 ± 0.16 | 2.43 ± 0.36 |  |  |  |
| Hematocrit<br>(%) | ♀ | 20.08 ± 2.55 | 23.77 ± 0.32 | 19.85 ± 4.01 | 0.351 | 0.173 | 0.751 |
|  | ♂ | 22.26 ± 2.48 | 24.83 ± 0.32 | 24.25 ± 2.06 |  |  |  |
| Hemoglobin<br>(g/dL) | ♀ | 7.82 ± 0.68 | 9.07 ± 0.15 | 8.43 ± 1.11 | 0.287 | 0.181 | 0.714 |
|  | ♂ | 8.62 ± 0.58 | 9.20 ± 0.12 | 9.49 ± 0.53 |  |  |  |
| MCV<br>(fL) | ♀ | 89.23 ± 1.05 | 92.87 ± 1.88 | 83.86 ± 5.52 | 0.092 | 0.789 | 0.943 |
|  | ♂ | 88.90 ± 1.92 | 90.70 ± 1.70 | 83.99 ± 5.86 |  |  |  |
| MCH<br>(pg) | ♀ | 37.05 ± 3.53 | 35.03 ± 0.86 | 71.20 ± 24.69 | 0.109 | 0.921 | 0.737 |
|  | ♂ | 35.96 ± 2.55 | 44.89 ± 11.01 | 58.91 ± 21.84 |  |  |  |
| MCHC<br>(g/L) | ♀ | 415.43 ± 39.78 | 377.44 ± 6.08 | 981.07 ± 309.80 | 0.100 | 0.928 | 0.938 |
|  | ♂ | 405.67 ± 29.86 | 505.69 ± 133.20 | 923.44 ± 508.45 |  |  |  |
| RDW-CV<br>(%) | ♀ | 29.11 ± 3.17 | 24.01 ± 1.37 | 34.18 ± 5.84 | 0.134 | 0.632 | 0.601 |
|  | ♂ | 30.12 ± 4.63 | 24.40 ± 2.11 | 28.42 ± 3.86 |  |  |  |
| RDW-SD<br>(fL) | ♀ | 106.46 ± 12.54 | 90.24 ± 6.70 | 114.49 ± 12.94 | 0.226 | 0.558 | 0.538 |
|  | ♂ | 110.97 ± 18.64 | 89.27 ± 8.76 | 94.36 ± 7.45 |  |  |  |
| Platelets<br>(10<sup>3</sup>/μL) | ♀ | 739.21 ± 196.91 | 534.84 ± 98.04 | 1121.07 ± 98.04 | 0.060 | 0.791 | 0.909 |
|  | ♂ | 853.91 ± 275.52 | 620.08 ± 71.52 | 1060.58 ± 71.52 |  |  |  |
| MPV<br>(fL) | ♀ | 8.61 ± 0.32 | 8.49 ± 0.32 | 8.57 ± 0.52 | 0.694 | 0.751 | 0.733 |
|  | ♂ | 8.81 ± 0.38 | 8.43 ± 0.40 | 8.12 ± 0.50 |  |  |  |
| PDW<br>(%) | ♀ | 18.58 ± 0.38 | 18.84 ± 0.43 | 18.17 ± 0.42 | 0.328 | 0.967 | 0.897 |
|  | ♂ | 18.38 ± 0.36 | 18.89 ± 0.42 | 18.35 ± 0.43 |  |  |  |
| PCT | ♀ | 0.47 ± 0.05 | 0.40 ± 0.07 | 0.70 ± 0.08 | 0.278 | 0.777 | 0.063 |

The results correspond to hematological parameters of female and male pups from sleep restricted or control dams. Data were obtained on postpartum day 9 and are presented as mean ± SEM. A two-way ANOVA followed by Tukey's *post hoc* test was used. The symbols \* and # indicate significant differences compared to the control group (\*, $p < 0.05$ ; \*\*, $p < 0.01$ ) and the ASR group (#, $p < 0.05$ ), respectively. ASR, acute sleep restriction; CSR, chronic sleep restriction; MCV, mean corpuscular volume; MCH, mean corpuscular hemoglobin; MCHC, mean corpuscular hemoglobin concentration; RDW-CV, red blood cell distribution width – coefficient of variation; RDW-SD, red blood cell distribution width – standard deviation; MPV, mean platelet volume; PDW, platelet distribution width; PCT, plateletcrit.

As an approach to examine whether lymphocyte counts in pups were influenced by the number of days of maternal sleep restriction, simple linear regression analyses were conducted. The results revealed a positive relationship between these two parameters for both females (slope = 0.3515, R² = 0.33, p = 0.005) and males (slope = 0.2582, R² = 0.34, p = 0.003).

Two-way ANOVA indicated a significant effect of experimental condition on total neutrophil counts in pups (p = 0.026), although *post hoc* tests did not reach significance. Other blood parameters showed no significant differences in either mothers or pups, and sex-based analyses of leukocytes were also non-significant (Table 2). The neutrophil-to-lymphocyte ratio, an indicator of stressLJ^74^, also showed no significant differences among control mothers (0.39 ± 0.09), ASR (0.55 ± 0.04), and CSR (0.36 ± 0.06; F = 2.33; p = 0.123). Similarly, no significant differences were observed either in females (C: 0.31 ± 0.05; ASR: 0.25 ± 0.04; CSR: 0.27 ± 0.03) or male pups (C: 0.29 ± 0.04; ASR: 0.25 ± 0.04; CSR: 0.26 ± 0.03), with no significant experimental condition (p = 0.444), sex (p = 0.753), or interaction effects (p = 0.973).

### 3.3. Variables correlations

Figure 3 shows the Spearman correlation matrices, which were used to evaluate the associations among the measured variables and their potential group-dependent variations.

**Figure 3.**
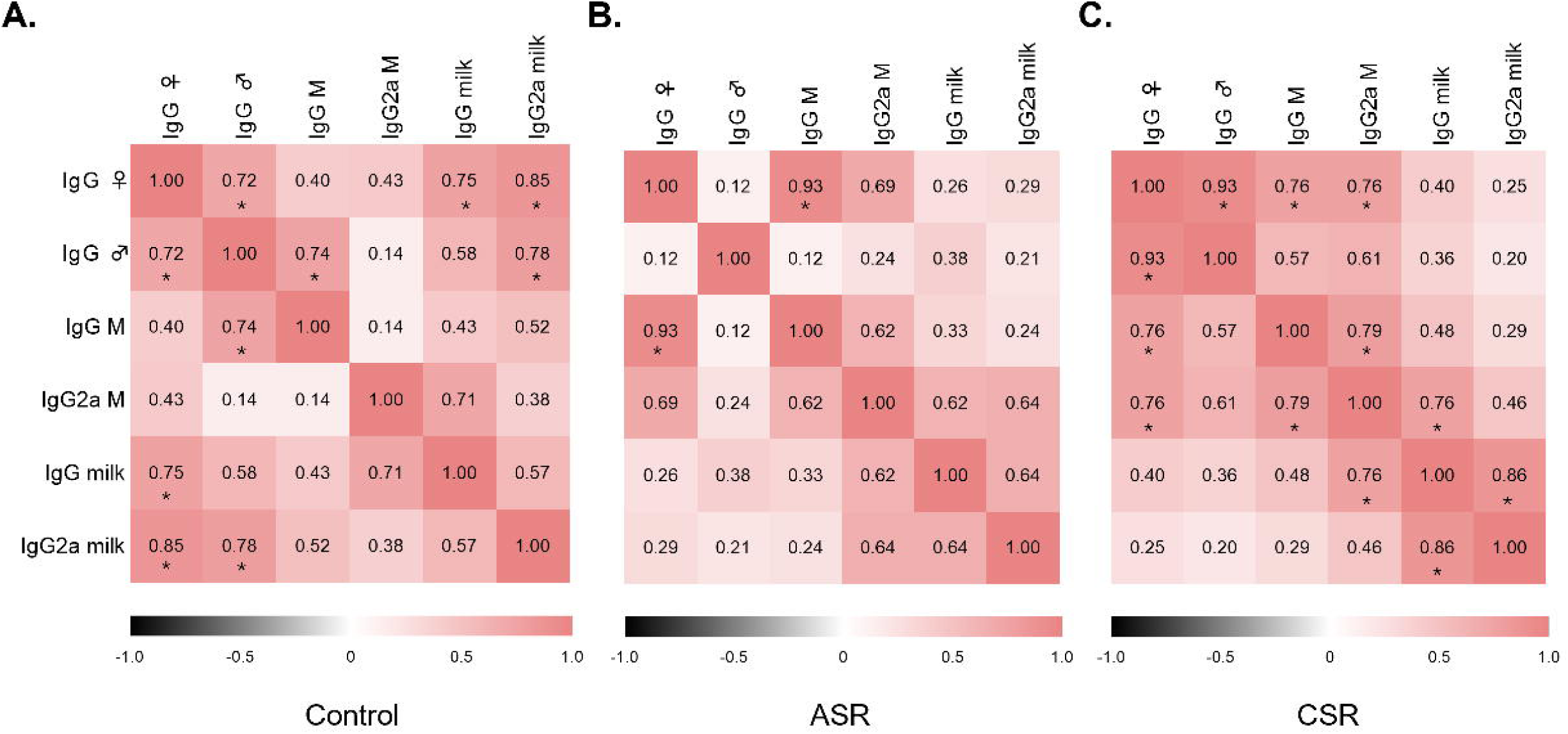
Humoral variable correlations. **A-C**. Heat maps representations of Spearman correlations matrix between the serum and milk parameters measured after control **(A)**, acute sleep restriction **(B)** or chronic sleep restriction **(C)** procedures. Black-grey boxes represent negative correlations (Spearman r < 0), white boxes correspond to no correlation (Spearman r = 0), and pink boxes denote a positive correlation (Spearman r > 0). The asterisk symbols (*) denote statistical significance (p < 0.05). IgG ♀, IgG levels in serum from female pups; IgG ♂, IgG levels in serum from female pups; IgG M, IgG serum maternal levels; IgG2a M, IgG2a serum maternal levels; IgG milk, IgG milk levels; IgG2a milk, IgG2a milk levels; ASR, acute sleep restriction; CSR, chronic sleep restriction.

Regarding the correlations between antibody levels in the pups and maternal serum, IgG levels were significantly correlated across all groups, with sex-specific differences in the pups. While in the control group the positive correlation was observed only in male pups (p = 0.046), in the sleep-restricted groups the correlation was found in female pups, both in the ASR (p = 0.002) and the CSR group (p = 0.037). Also, IgG levels from female and male pups were correlated in the control (p = 0.037) and CSR (p = 0.007) groups, but not in the ASR group (p = 0.793).

As for the correlations between IgG concentration in the serum of the pups and maternal milk antibodies, we observed a positive correlation with milk IgG in females (p = 0.025) and with milk IgG2a in both males (p = 0.017) and females (p = 0.006), but only in the control group.

## 4. Discussion

In the present study, we show the immunological effects of acute and chronic maternal sleep restriction on both dams and their pups in rats. Our main findings reveal that, although maternal sleep restriction did not affect any of the maternal immunological parameters measured, it altered serum antibody levels and leukocyte counts in the offspring. Specifically, we found lower IgG serum levels in male pups from mothers subjected to acute sleep restriction, and higher blood lymphocytes counts in both female and male offspring of mothers exposed to chronic sleep restriction.

### 4.1. Maternal immune parameters were unaffected by sleep restriction

Neither ASR nor CSR affected the measured maternal immunological parameters. Specifically, milk and serum IgG and IgG2a, serum IL-17A and IL-6, and hematological parameters remained unchanged. Previous works, in stages outside the postpartum period, showed that these parameters could be affected by sleep disturbances, depending on the methodology and duration of sleep deprivation ^32–35,75–78^. Thus, the seemingly contradictory results with literature may be attributed, on one hand, to the protocol employed. Specifically, six hours of sleep restriction for one or five days, with an 18-hour recovery period between sessions during chronic restriction, might be too subtle to induce significant effects on the immune system. In this context, we emphasize the importance of investigating milder sleep restriction models that more accurately reflect typical human conditions, where prolonged total sleep deprivation is uncommon.

On the other hand, we hypothesize that the lack of observed changes may be related to the unique physiology of the postpartum period ^96–98^, during which mothers might adapt to additional sleep disruption by exhibiting either a higher threshold for immunological alterations following sleep deprivation or a capacity for rapid recovery during the 18-hour recovery intervals. Considering that sleep disturbances are an intrinsic feature of motherhood, a persistent proinflammatory state induced by sleep deprivation would likely be detrimental to both the mother and the offspring ^52^.

Regarding changes in immunological parameters due to sleep alterations during the postpartum period, studies have shown no significant differences in IL-6 concentrations between women with and without insomnia ^52^. However, sleeping five or fewer hours per day or experiencing lower sleep quality has been associated with higher levels of this cytokine ^53,79^. Gröer and colleagues ^50^ reported that mothers with higher levels of sleepiness and fatigue presented more symptoms of infection. Nevertheless, no direct reports exist regarding changes in the quantity or proportion of white blood cells in response to sleep deprivation. To our knowledge, this is the first study to examine the effects of sleep loss on both antibody concentrations and white blood cell counts in postpartum mothers.

### 4.2. Maternal sleep restriction impaired immunoglobulin concentrations and lymphocyte count in the offspring

Despite the absence of changes in maternal parameters, certain immunological parameters from the pups were affected by maternal sleep restriction. Regarding IgG concentration, to our knowledge, no previous reports have documented changes in antibody levels in the offspring of sleep-deprived mothers. Given that sleep deprivation is inherently stressful ^2,80,81^, and that maternal stress exerts diverse effects on various offspring parameters ^82–86^, it would be expected that maternal sleep restriction could influence immunological processes of the pups. Consistent with our findings, maternal stress during the prenatal period has been shown to reduce IgG levels in the offspring ^87^.

Interestingly, the positive correlation between milk IgG and offspring serum IgG observed in the control group, was no longer present in the sleep restricted groups, suggesting that sleep deprivation may impair the transfer of IgG from milk to blood. Given that at postnatal day 9 serum IgG is largely milk-derived ^88^ and milk protein content is reduced in acutely sleep-deprived mothers ^60^, we hypothesize that the reduction in milk protein amount may reflect a change in transferred immunity that causes this pups’ IgG reduction. Although we did not observe a decrease in IgG levels in milk, variations in other bioactive proteins that regulate IgG concentrations in the offspring may mediate this effect, for example by influencing the transport of IgG from milk to blood across the intestinal wall ^89^.

The reduction in IgG due to maternal sleep restriction was not observed in female pups, suggesting that male offspring are more susceptible to maternal sleep deprivation, which is consistent with previous reports showing sex-based differences in offspring immune responses ^90–94^. One factor accounting for observed changes could be sex-related genetic differences associated with antibody uptake ^95–97^. In addition, mother rats behave differently toward male and female pups, for example, spending more time licking male pups ^98,99^, and male pups are more affected by alterations in maternal licking behavior ^100^. Although we did not analyze these behaviors separately by pup sex, we previously evidenced a reduction in maternal behaviors following sleep restriction ^60,64^. Thus, it is possible that the decrease in maternal licking was more pronounced in male pups, ultimately affecting them more.

In regard to cellular immunity, both male and female pups of mothers subjected to chronic sleep restriction exhibited a higher lymphocyte counts compared with the control group. In addition, we found a significant positive linear relationship between the number of days of maternal sleep deprivation and lymphocyte counts in the offspring, suggesting a cumulative impact of maternal sleep restriction on the pups’ immune function. To our knowledge, there is no previous literature demonstrating the effects of maternal sleep deprivation during either the prenatal or postnatal period on hematological parameters in offspring. Only one study, using self-reported questionnaires, showed that greater maternal sleepiness and fatigue were associated with increased symptoms of infection in both mothers and their children^50^.

Regarding the causes of the rise observed in lymphocyte count in pups, changes in maternal–pup interactions, such as maternal separation, have been shown to alter the offspring’s immune system ^101^. In this context, in the cohort of animals from the present study, we observed a decrease in licking behaviors following maternal chronic sleep restriction ^60^. This leads us to hypothesize that alterations in maternal behavior may contribute to the elevated lymphocyte counts observed in the pups.

Another factor contributing to the increased lymphocyte count in pups is that chronic maternal sleep restriction could disrupt lymphocyte migration in the offspring via milk-transferred mediators ^102,103^. Sleep deprivation impairs the expression of lymphocyte adhesion molecules involved in tissue migration, reducing migratory capacity and increasing their concentration in the blood ^49,104^. Since this effect is regulated by growth hormone and prolactin ^49^, bioactive mediators such as prolactin or growth hormone transferred through milk ^105,106^ could contribute to the observed increase in lymphocyte counts. Further studies are needed to clarify how maternal sleep deprivation affects the offspring’s immune system.

### 4.3. Technical considerations

An important consideration is the potential impact of anesthesia, surgery and the timing of sample collection on humoral and cellular immune parameters. However, any differences observed between groups cannot be attributed to these factors, since the anesthesia doses and timing protocol were identical for all mothers across the three experimental groups.

Finally, although the sample size per variable in the statistical analysis of the Spearman correlation matrices and linear regression was low for extracting consistent conclusions, the resulting data provide valuable insights that support the formulation of new hypotheses.

### 4.4. Final remarks

Maternal sleep restriction altered serum antibody and leukocyte levels in the offspring without affecting maternal immune parameters, suggesting that, even in the absence of detectable maternal immune changes, the offspring are impacted, with male pups being particularly susceptible. These immunological effects restricted to the pups suggests the involvement of bioactive mediators between mothers and pups, potentially transmitted through milk or either caused by the deterioration of maternal behavior previously shown in these rats ^60^. Further investigations are required to determine the underlying cause of these findings. Given the scarce literature on immunological parameters in mother-offspring dyads following sleep deprivation, and the limited research on the impact of sleep deprivation on immune function in females of any species, we consider our work to be of significant value and emphasizes the need for further studies in this area.

## Author Contributions

The authors’ contributions were as follows: conceptualization: L.B.; resources: A.H., A.S., P.T., T.F., L.B.; investigation: F.P., C.R-C., A.H., M.R., A.S., D.S., J.P.C, T.F. and L.B.; formal analysis: F.P.; supervision: A.H. and L.B.; methodology: L.B. and F.P.; writing – original draft: F.P., A.H. and L.B.; writing – review & editing: F.P., C.R-C., A.H., M.R., A.S., D.S., J.P.C., P.T., T.F. and L.B.; project administration: L.B.; funding acquisition: L.B. and F.P. All authors have read and agreed to the published version of the manuscript.

## Institutional Review Board Statement

All the experimental procedures were approved by the Institutional Animal Care Committee (protocol number 070151-000008-22) and were undertaken in conformity with the “Guide to the Care and Use of Laboratory Animals” (8th edition, National Academy Press, Washington D.C., 2011).

## Informed Consent Statement

Not applicable.

## Data Availability Statement

Not applicable.

## References

1. Belenky G, Wesensten NJ, Thorne DR, et al. Patterns of performance degradation and restoration during sleep restriction and subsequent recovery: a sleep dose-response study. J Sleep Res. 2003;12(1):1–12. doi:10.1046/j.1365-2869.2003.00337.x

2. Meerlo P, Koehl M, van der Borght K, Turek FW. Sleep restriction alters the hypothalamic-pituitary-adrenal response to stress. J Neuroendocrinol. 2002;14(5):397–402. doi:10.1046/j.0007-1331.2002.00790.x

3. Meerlo P, Sgoifo A, Suchecki D. Restricted and disrupted sleep: effects on autonomic function, neuroendocrine stress systems and stress responsivity. Sleep Med Rev. 2008;12(3):197–210. doi:10.1016/j.smrv.2007.07.007

4. Besedovsky L, Lange T, Haack M. The Sleep-Immune Crosstalk in Health and Disease. Physiol Rev. 2019;99(3):1325–1380. doi:10.1152/physrev.00010.2018

5. Sivertsen B, Hysing M, Dørheim SK, Eberhard-Gran M. Trajectories of maternal sleep problems before and after childbirth: a longitudinal population-based study. BMC Pregnancy Childbirth. 2015;15:129. doi:10.1186/s12884-015-0577-1

6. Creti L, Libman E, Rizzo D, et al. Sleep in the Postpartum: Characteristics of First-Time, Healthy Mothers. Sleep Disord. 2017;2017:8520358. doi:10.1155/2017/8520358

7. Okun ML, Lac A. Postpartum Insomnia and Poor Sleep Quality Are Longitudinally Predictive of Postpartum Mood Symptoms. Psychosom Med. 2023;85(8):736–743. doi:10.1097/PSY.0000000000001234

8. Manconi M, van der Gaag LC, Mangili F, et al. Sleep and sleep disorders during pregnancy and postpartum: The Life-ON study. Sleep Med. 2024;113:41–48. doi:10.1016/j.sleep.2023.10.021

9. Bei B, Coo S, Trinder J. Sleep and Mood During Pregnancy and the Postpartum Period. Sleep Med Clin. 2015;10(1):25–33. doi:10.1016/j.jsmc.2014.11.011

10. Hunter LP, Rychnovsky JD, Yount SM. A selective review of maternal sleep characteristics in the postpartum period. J Obstet Gynecol Neonatal Nurs JOGNN. 2009;38(1):60–68. doi:10.1111/j.1552-6909.2008.00309.x

11. Lyamin O, Pryaslova J, Kosenko P, Siegel J. Behavioral aspects of sleep in bottlenose dolphin mothers and their calves. Physiol Behav. 2007;92(4):725–733. doi:10.1016/j.physbeh.2007.05.064

12. Sivadas N, Radhakrishnan A, Aswathy BS, Kumar VM, Gulia KK. Dynamic changes in sleep pattern during post-partum in normal pregnancy in rat model. Behav Brain Res. 2017;320:264–274. doi:10.1016/j.bbr.2016.11.040

13. Benedetto L, Rivas M, Pereira M, Ferreira A, Torterolo P. A descriptive analysis of sleep and wakefulness states during maternal behaviors in postpartum rats. Arch Ital Biol. 2017;155(3):99–109. doi:10.12871/00039829201731

14. Benedetto L, Peña F, Rivas M, Ferreira A, Torterolo P. The Integration of the Maternal Care with Sleep During the Postpartum Period. Sleep Med Clin. 2023;18(4):499–509. doi:10.1016/j.jsmc.2023.06.013

15. Komiya H, Miyoshi C, Iwasaki K, et al. Sleep/Wake Behaviors in Mice During Pregnancy and Pregnancy-Associated Hypertensive Mice. Sleep. 2018;41(3):zsx209. doi:10.1093/sleep/zsx209

16. Tucker P, Leineweber C, Kecklund G. Comparing the acute effects of shiftwork on mothers and fathers. Occup Med Oxf Engl. 2021;71(9):414–421. doi:10.1093/occmed/kqab083

17. Kecklund G, Axelsson J. Health consequences of shift work and insufficient sleep. BMJ. 2016;355:i5210. doi:10.1136/bmj.i5210

18. Dørheim SK, Bondevik GT, Eberhard-Gran M, Bjorvatn B. Sleep and Depression in Postpartum Women: A Population-Based Study. Sleep. 2009;32(7):847–855.

19. Kalil A, Dunifon R, Crosby D, Su JH. Work Hours, Schedules, and Insufficient Sleep Among Mothers and Their Young Children. J Marriage Fam. 2014;76(5):891–904. doi:10.1111/jomf.12142

20. Moldofsky H. Central nervous system and peripheral immune functions and the sleep-wake system. J Psychiatry Neurosci JPN. 1994;19(5):368–374.

21. Opp MR, Imeri L. Sleep as a behavioral model of neuro-immune interactions. Acta Neurobiol Exp (Warsz). 1999;59(1):45–53. doi:10.55782/ane-1999-1295

22. Irwin M. Effects of sleep and sleep loss on immunity and cytokines. Brain Behav Immun. 2002;16(5):503–512. doi:10.1016/s0889-1591(02)00003-x

23. Irwin MR, Olmstead R, Carroll JE. Sleep Disturbance, Sleep Duration, and Inflammation: A Systematic Review and Meta-Analysis of Cohort Studies and Experimental Sleep Deprivation. Biol Psychiatry. 2016;80(1):40–52. doi:10.1016/j.biopsych.2015.05.014

24. Krueger JM, Majde JA. Cytokines and sleep. Int Arch Allergy Immunol. 1995;106(2):97–100. doi:10.1159/000236827

25. Besedovsky L, Lange T, Born J. Sleep and immune function. Pflüg Arch – Eur J Physiol. 2012;463(1):121–137. doi:10.1007/s00424-011-1044-0

26. Krueger JM, Walter J, Dinarello CA, Wolff SM, Chedid L. Sleep-promoting effects of endogenous pyrogen (interleukin-1). Am J Physiol. 1984;246(6 Pt 2):R994–999. doi:10.1152/ajpregu.1984.246.6.R994

27. Tobler I, Borbély AA, Schwyzer M, Fontana A. Interleukin-1 derived from astrocytes enhances slow wave activity in sleep EEG of the rat. Eur J Pharmacol. 1984;104(1-2):191–192. doi:10.1016/0014-2999(84)90391-1

28. Opp MR, Obal F, Krueger JM. Interleukin 1 alters rat sleep: temporal and dose-related effects. Am J Physiol. 1991;260(1 Pt 2):R52–58. doi:10.1152/ajpregu.1991.260.1.R52

29. Vgontzas AN, Zoumakis E, Bixler EO, et al. Adverse effects of modest sleep restriction on sleepiness, performance, and inflammatory cytokines. J Clin Endocrinol Metab. 2004;89(5):2119–2126. doi:10.1210/jc.2003-031562

30. Taishi P, Chen Z, Obál F, et al. Sleep-associated changes in interleukin-1beta mRNA in the brain. J Interferon Cytokine Res Off J Int Soc Interferon Cytokine Res. 1998;18(9):793–798. doi:10.1089/jir.1998.18.793

31. van Leeuwen WMA, Lehto M, Karisola P, et al. Sleep restriction increases the risk of developing cardiovascular diseases by augmenting proinflammatory responses through IL-17 and CRP. PloS One. 2009;4(2):e4589. doi:10.1371/journal.pone.0004589

32. Yehuda S, Sredni B, Carasso RL, Kenigsbuch-Sredni D. REM sleep deprivation in rats results in inflammation and interleukin-17 elevation. J Interferon Cytokine Res Off J Int Soc Interferon Cytokine Res. 2009;29(7):393–398. doi:10.1089/jir.2008.0080

33. Sang D, Lin K, Yang Y, et al. Prolonged sleep deprivation induces a cytokine-storm-like syndrome in mammals. Cell. 2023;186(25):5500–5516.e21. doi:10.1016/j.cell.2023.10.025

34. Everson CA. Clinical assessment of blood leukocytes, serum cytokines, and serum immunoglobulins as responses to sleep deprivation in laboratory rats. Am J Physiol-Regul Integr Comp Physiol. 2005;289(4):R1054–R1063. doi:10.1152/ajpregu.00021.2005

35. Hui L, Hua F, Diandong H, Hong Y. Effects of sleep and sleep deprivation on immunoglobulins and complement in humans. Brain Behav Immun. 2007;21(3):308–310. doi:10.1016/j.bbi.2006.09.005

36. Yang SW, Yang HF, Chen YY, Chen WL. Sleep deprivation and immunoglobulin E level. Front Med. 2022;9. doi:10.3389/fmed.2022.955085

37. Renegar KB, Floyd R, Krueger JM. Effect of Sleep Deprivation on Serum Influenza-Specific IgG. Sleep. 1998;21(1):19–24. doi:10.1093/sleep/21.1.19

38. Lahimgarzadeh R, Vaseghi S, Nasehi M, Rouhollah F. Effect of multi-epitope derived from HIV-1 on REM sleep deprivation-induced spatial memory impairment with respect to the level of immune factors in mice. Iran J Basic Med Sci. 2022;25(2):164–172. doi:10.22038/IJBMS.2022.61175.13536

39. Spiegel K, Rey AE, Cheylus A, et al. A meta-analysis of the associations between insufficient sleep duration and antibody response to vaccination. Curr Biol. 2023;33(5):998–1005.e2. doi:10.1016/j.cub.2023.02.017

40. Rayatdoost E, Rahmanian M, Sanie MS, et al. Sufficient Sleep, Time of Vaccination, and Vaccine Efficacy: A Systematic Review of the Current Evidence and a Proposal for COVID-19 Vaccination. Yale J Biol Med. 2022;95(2):221–235.

41. Barriga-Ibars C, Rodríguez-Moratinos AB, Esteban S, Rial RV. Interrelations between sleep and the immune status. Rev Neurol. Invalid date;40(9):548–556. doi:10.33588/rn.4009.2005250

42. Hayashi M, Shimba S, Tezuka M. Characterization of the molecular clock in mouse peritoneal macrophages. Biol Pharm Bull. 2007;30(4):621–626. doi:10.1248/bpb.30.621

43. Irwin M, McClintick J, Costlow C, Fortner M, White J, Gillin JC. Partial night sleep deprivation reduces natural killer and cellular immune responses in humans. FASEB J Off Publ Fed Am Soc Exp Biol. 1996;10(5):643–653. doi:10.1096/fasebj.10.5.8621064

44. Hahn J, Günter M, Schuhmacher J, et al. Sleep enhances numbers and function of monocytes and improves bacterial infection outcome in mice. Brain Behav Immun. 2020;87:329–338. doi:10.1016/j.bbi.2020.01.001

45. Irwin M, Mascovich A, Gillin JC, Willoughby R, Pike J, Smith TL. Partial sleep deprivation reduces natural killer cell activity in humans. Psychosom Med. 1994;56(6):493–498. doi:10.1097/00006842-199411000-00004

46. Moldofsky H, Lue FA, Davidson JR, Gorczynski R. Effects of sleep deprivation on human immune functions. FASEB J Off Publ Fed Am Soc Exp Biol. 1989;3(8):1972–1977. doi:10.1096/fasebj.3.8.2785942

47. Ingram LA, Simpson RJ, Malone E, Florida-James GD. Sleep disruption and its effect on lymphocyte redeployment following an acute bout of exercise. Brain Behav Immun. 2015;47:100–108. doi:10.1016/j.bbi.2014.12.018

48. Said EA, Al-Abri MA, Al-Saidi I, et al. Sleep deprivation alters neutrophil functions and levels of Th1-related chemokines and CD4+ T cells in the blood. Sleep Breath Schlaf Atm. 2019;23(4):1331–1339. doi:10.1007/s11325-019-01851-1

49. Martínez-Albert E, Lutz ND, Hübener R, et al. Sleep promotes T-cell migration towards CCL19 via growth hormone and prolactin signaling in humans. Brain Behav Immun. 2024;118:69–77. doi:10.1016/j.bbi.2024.02.021

50. Groër M, Davis M, Casey K, Short B, Smith K, Groër S. Neuroendocrine and immune relationships in postpartum fatigue. MCN Am J Matern Child Nurs. 2005;30(2):133–138. doi:10.1097/00005721-200503000-00012

51. Corwin EJ, Bozoky I, Pugh LC, Johnston N. Interleukin-1beta elevation during the postpartum period. Ann Behav Med Publ Soc Behav Med. 2003;25(1):41–47. doi:10.1207/S15324796ABM2501_06

52. Drozdowicz-Jastrzębska E, Mach A, Skalski M, et al. Depression, anxiety, insomnia and interleukins in the early postpartum period. Front Psychiatry. 2023;14. doi:10.3389/fpsyt.2023.1266390

53. Taveras EM, Rifas-Shiman SL, Rich-Edwards JW, Mantzoros CS. Maternal short sleep duration is associated with increased levels of inflammatory markers at 3 years postpartum. Metabolism. 2011;60(7):982–986. doi:10.1016/j.metabol.2010.09.008

54. Proudfoot KL, Kull JA, Krawczel PD, et al. Effects of acute lying and sleep deprivation on metabolic and inflammatory responses of lactating dairy cows. J Dairy Sci. 2021;104(4):4764–4774. doi:10.3168/jds.2020-19332

55. Caldji C, Tannenbaum B, Sharma S, Francis D, Plotsky PM, Meaney MJ. Maternal care during infancy regulates the development of neural systems mediating the expression of fearfulness in the rat. Proc Natl Acad Sci U S A. 1998;95(9):5335–5340. doi:10.1073/pnas.95.9.5335

56. Meaney MJ, Szyf M. Environmental programming of stress responses through DNA methylation: life at the interface between a dynamic environment and a fixed genome. Dialogues Clin Neurosci. 2005;7(2):103–123.

57. Gomez de Agüero M, Ganal-Vonarburg SC, Fuhrer T, et al. The maternal microbiota drives early postnatal innate immune development. Science. 2016;351(6279):1296–1302. doi:10.1126/science.aad2571

58. Cebra JJ. Influences of microbiota on intestinal immune system development2. Am J Clin Nutr. 1999;69(5):1046S–1051S. doi:10.1093/ajcn/69.5.1046s

59. Trim MJ, Wheeler RV, Franklin TB. Maternal immune activation accelerates pup reflex development and alters immune proteins in pup stomach contents and brain. Brain Res. 2024;1845:149198. doi:10.1016/j.brainres.2024.149198

60. Peña F, Serantes D, Rivas M, et al. Acute and chronic sleep restriction differentially modify maternal behavior and milk macronutrient composition in the postpartum rat. Physiol Behav. 2024;278:114522. doi:10.1016/j.physbeh.2024.114522

61. National Research Council (US) Committee for the Update of the Guide for theCare and Use of Laboratory Animals. Guide for the Care and Use of Laboratory Animals. 8th ed. National Academies Press (US); 2011. Accessed June 6, 2025. http://www.ncbi.nlm.nih.gov/books/NBK54050/

62. Benedetto L, Rivas M, Cavelli M, et al. Microinjection of the dopamine D2-receptor antagonist Raclopride into the medial preoptic area reduces REM sleep in lactating rats. Neurosci Lett. 2017;659:104–109. doi:10.1016/j.neulet.2017.08.077

63. Peña F, Benedetto L, Rivas M, et al. Sleep and maternal behavior in the postpartum rat after haloperidol and midazolam treatments SPECIAL ISSUE. Sleep Sci. Published online July 27, 2020. doi:10.5935/1984-0063.20200019

64. Peña F, Rivas M, Serantes D, Ferreira A, Torterolo P, Benedetto L. Is sleep critical for lactation in rat? Physiol Behav. 2022;258:114011. doi:10.1016/j.physbeh.2022.114011

65. Paxinos G, Watson C. The Rat Brain in Stereotaxic Coordinates–The New Coronal Set, 5th Edn.; 2004.

66. Benedetto L, Rivas M, Peña F, Serantes D, Ferreira A, Torterolo P. Local administration of bicuculline into the ventrolateral and medial preoptic nuclei modifies sleep and maternal behavior in lactating rats. Physiol Behav. 2021;238:113491. doi:10.1016/j.physbeh.2021.113491

67. Rivas M, Serantes D, Peña F, et al. Role of Hypocretin in the Medial Preoptic Area in the Regulation of Sleep, Maternal Behavior and Body Temperature of Lactating Rats. Neuroscience. 2021;475:148–162. doi:10.1016/j.neuroscience.2021.08.034

68. DePeters EJ, Hovey RC. Methods for collecting milk from mice. J Mammary Gland Biol Neoplasia. 2009;14(4):397–400. doi:10.1007/s10911-009-9158-0

69. Rodgers CT. Practical aspects of milk collection in the rat. Lab Anim. 1995;29(4):450–455. doi:10.1258/002367795780739980

70. Muranishi Y, Parry L, Averous J, et al. Method for collecting mouse milk without exogenous oxytocin stimulation. BioTechniques. 2016;60(1):47–49. doi:10.2144/000114373

71. Benedetto L, Rodriguez-Servetti Z, Lagos P, D’Almeida V, Monti J, Torterolo P. Microinjection of Melanin Concentrating Hormone into the lateral preoptic area promotes non-REM sleep in the rat. Peptides. 2012;39. doi:10.1016/j.peptides.2012.10.005

72. Serantes D, Cavelli M, Gonzalez J, Mondino A, Benedetto L, Torterolo P. Characterising the power spectrum dynamics of the non-REM to REM sleep transition. J Sleep Res. 2025;34(3):e14388. doi:10.1111/jsr.14388

73. Sánchez MB, Michel Lara MC, Neira FJ, et al. Hyperthyroidism keeps immunoglobulin levels but reduces milk fat and CD11b/c+ cells on early lactation. Mol Cell Endocrinol. 2024;594:112370. doi:10.1016/j.mce.2024.112370

74. Swan MP, Hickman DL. Evaluation of the neutrophil-lymphocyte ratio as a measure of distress in rats. Lab Anim. 2014;43(8):276–282. doi:10.1038/laban.529

75. Hu J, Chen Z, Gorczynski CP, et al. Sleep-deprived mice show altered cytokine production manifest by perturbations in serum IL-1ra, TNFa, and IL-6 levels. Brain Behav Immun. 2003;17(6):498–504. doi:10.1016/j.bbi.2003.03.001

76. Boudjeltia KZ, Faraut B, Stenuit P, et al. Sleep restriction increases white blood cells, mainly neutrophil count, in young healthy men: A pilot study. Vasc Health Risk Manag. 2008;4(6):1467–1470.

77. Lasselin J, Rehman JU, Åkerstedt T, Lekander M, Axelsson J. Effect of long-term sleep restriction and subsequent recovery sleep on the diurnal rhythms of white blood cell subpopulations. Brain Behav Immun. 2015;47:93–99. doi:10.1016/j.bbi.2014.10.004

78. Zager A, Andersen ML, Ruiz FS, Antunes IB, Tufik S. Effects of acute and chronic sleep loss on immune modulation of rats. Am J Physiol-Regul Integr Comp Physiol. 2007;293(1):R504–R509. doi:10.1152/ajpregu.00105.2007

79. Christian LM, Kowalsky JM, Mitchell AM, Porter K. Associations of postpartum sleep, stress, and depressive symptoms with LPS-stimulated cytokine production among African American and White women. J Neuroimmunol. 2018;316:98–106. doi:10.1016/j.jneuroim.2017.12.020

80. Mirescu C, Peters JD, Noiman L, Gould E. Sleep deprivation inhibits adult neurogenesis in the hippocampus by elevating glucocorticoids. Proc Natl Acad Sci U S A. 2006;103(50):19170–19175. doi:10.1073/pnas.0608644103

81. Pires GN, Bezerra AG, Tufik S, Andersen ML. Effects of experimental sleep deprivation on anxiety-like behavior in animal research: Systematic review and meta-analysis. Neurosci Biobehav Rev. 2016;68:575–589. doi:10.1016/j.neubiorev.2016.06.028

82. Götz AA, Stefanski V. Psychosocial maternal stress during pregnancy affects serum corticosterone, blood immune parameters and anxiety behaviour in adult male rat offspring. Physiol Behav. 2007;90(1):108–115. doi:10.1016/j.physbeh.2006.09.014

83. Llorente E, Brito ML, Machado P, González MC. Effect of prenatal stress on the hormonal response to acute and chronic stress and on immune parameters in the offspring. J Physiol Biochem. 2002;58(3):143–149. doi:10.1007/BF03179851

84. Catalani A, Casolini P, Cigliana G, et al. Maternal corticosterone influences behavior, stress response and corticosteroid receptors in the female rat. Pharmacol Biochem Behav. 2002;73(1):105–114. doi:10.1016/s0091-3057(02)00755-4

85. Hayashi A, Nagaoka M, Yamada K, Ichitani Y, Miake Y, Okado N. Maternal stress induces synaptic loss and developmental disabilities of offspring. Int J Dev Neurosci. 1998;16(3):209–216. doi:10.1016/S0736-5748(98)00028-8

86. Galley JD, Mashburn-Warren L, Blalock LC, et al. Maternal anxiety, depression and stress affects offspring gut microbiome diversity and bifidobacterial abundances. Brain Behav Immun. 2023;107:253–264. doi:10.1016/j.bbi.2022.10.005

87. Sobrian SK, Vaughn VT, Bloch EF, Burton LE. Influence of prenatal maternal stress on the immunocompetence of the offspring. Pharmacol Biochem Behav. 1992;43(2):537–547. doi:10.1016/0091-3057(92)90189-M

88. Sewald K, Mueller M, Buschmann J, Hansen T, Lewin G. Development of hematological and immunological characteristics in neonatal rats. Reprod Toxicol. 2015;56:109–117. doi:10.1016/j.reprotox.2015.05.019

89. Jiang L, Wang J, Solorzano-Vargas RS, et al. Characterization of the rat intestinal Fc receptor (FcRn) promoter: transcriptional regulation of FcRn gene by the Sp family of transcription factors. Am J Physiol-Gastrointest Liver Physiol. 2004;286(6):G922–G931. doi:10.1152/ajpgi.00131.2003

90. Yoshimura S, Sakamoto S, Kudo H, Sassa S, Kumai A, Okamoto R. Sex-differences in adrenocortical responsiveness during development in rats. Steroids. 2003;68(5):439–445. doi:10.1016/S0039-128X(03)00045-X

91. Kühnemann S, Brown TJ, Hochberg RB, MacLusky NJ. Sex Differences in the Development of Estrogen Receptors in the Rat Brain. Horm Behav. 1994;28(4):483–491. doi:10.1006/hbeh.1994.1046

92. Rambousek L, Kacer P, Syslova K, Bumba J, Bubenikova-Valesova V, Slamberova R. Sex differences in methamphetamine pharmacokinetics in adult rats and its transfer to pups through the placental membrane and breast milk. Drug Alcohol Depend. 2014;139:138–144. doi:10.1016/j.drugalcdep.2014.03.023

93. Kosyreva AM, Dzhalilova DS, Makarova OV, et al. Sex differences of inflammatory and immune response in pups of Wistar rats with SIRS. Sci Rep. 2020;10(1):15884. doi:10.1038/s41598-020-72537-y

94. Klein SL, Flanagan KL. Sex differences in immune responses. Nat Rev Immunol. 2016;16(10):626–638. doi:10.1038/nri.2016.90

95. Kuilamu E, Subasic C, Cowin GJ, Simpson F, Minchin RF, Kaminskas LM. Cetuximab Exhibits Sex Differences in Lymphatic Exposure after Intravenous Administration in Rats in the Absence of Differences in Plasma Exposure. Pharm Res. 2020;37(11):224. doi:10.1007/s11095-020-02945-2

96. Surman SL, Jones BG, Penkert RR, et al. How Estrogen, Testosterone, and Sex Differences Influence Serum Immunoglobulin Isotype Patterns in Mice and Humans. Viruses. 2023;15(2):482. doi:10.3390/v15020482

97. Coutinho E, Jacobson L, Shock A, Smith B, Vernon A, Vincent A. Inhibition of Maternal-to-Fetal Transfer of IgG Antibodies by FcRn Blockade in a Mouse Model of Arthrogryposis Multiplex Congenita. Neurol Neuroimmunol Neuroinflammation. 2021;8(4):e1011. doi:10.1212/NXI.0000000000001011

98. Moore CL, Morelli GA. Mother rats interact differently with male and female offspring. J Comp Physiol Psychol. 1979;93(4):677–684. doi:10.1037/h0077599

99. Richmond G, Sachs BD. Maternal discrimination of pup sex in rats. Dev Psychobiol. 1984;17(1):87–89. doi:10.1002/dev.420170108

100. Reis AR, de Azevedo MS, de Souza MA, et al. Neonatal handling alters the structure of maternal behavior and affects mother–pup bonding. Behav Brain Res. 2014;265:216–228. doi:10.1016/j.bbr.2014.02.036

101. Roque S, Mesquita AR, Palha JA, Sousa N, Correia-Neves M. The Behavioral and Immunological Impact of Maternal Separation: A Matter of Timing. Front Behav Neurosci. 2014;8. doi:10.3389/fnbeh.2014.00192

102. Torres-Castro P, Abril-Gil M, Rodríguez-Lagunas MJ, Castell M, Pérez-Cano FJ, Franch À. TGF-β2, EGF, and FGF21 Growth Factors Present in Breast Milk Promote Mesenteric Lymph Node Lymphocytes Maturation in Suckling Rats. Nutrients. 2018;10(9):1171. doi:10.3390/nu10091171

103. Grases-Pintó B, Abril-Gil M, Rodríguez-Lagunas MJ, Castell M, Pérez-Cano FJ, Franch À. Leptin and adiponectin supplementation modifies mesenteric lymph node lymphocyte composition and functionality in suckling rats. Br J Nutr. 2018;119(5):486–495. doi:10.1017/S0007114517003786

104. Redwine L, Dang J, Irwin M. Cellular adhesion molecule expression, nocturnal sleep, and partial night sleep deprivation. Brain Behav Immun. 2004;18(4):333–340. doi:10.1016/j.bbi.2004.01.001

105. Whitworth NS, Grosvenor CE. Transfer of milk prolactin to the plasma of neonatal rats by intestinal absorption. J Endocrinol. 1978;79(2):191–199. doi:10.1677/joe.0.0790191

106. Grosvenor CE, Picciano MF, Baumrucker CR. Hormones and Growth Factors in Milk. Endocr Rev. 1993;14(6):710–728. doi:10.1210/edrv-14-6-710

